# A Mathematical Model to Predict Instantaneous Bone Formation Rate from Temporal Data of Cellular Biomarkers

**DOI:** 10.1101/2025.09.13.676058

**Authors:** Achsah Marlene Aruva, Jitendra Prasad

## Abstract

Although several models have been proposed to predict spatial patterns of new bone formation across different regions of a bone, to our knowledge no model has predicted bone formation temporally from cellular biomarkers. In this article, we predict bone formation rate (BFR) temporally from Col1a1 gene expression data and compare our predictions with the average BFR reported in the literature. The proposed mathematical model identifies key parameters influencing BFR at the celluar level and quantifies how biomarkers encode mechanical loading information. This model serves as an excellent starting point to understand how BFR changes over time relative to a given regimen of exogenous loading. We report that the simplest mathematical model explaining this phenonmenon with reasonable accuracy is a second order linear critically-damped system with a delay time. Since BFR reported in the literature typically represents an average value over the interlabel period, we also propose a method to convert this measure into an instantaneous one, which is essential for constructing an “ideal” mathematical model. Finally, we present our results and discuss limitations of the model along with directions for future improvement.

## 1. Introduction

The skeletal system is dynamic and adapts continuously in response to various biological and physical stimuli [1]. Numerous studies have demonstrated that mechanical loading profoundly influences bone’s structural adaptation [2, 3]. Bone contains an extensive network of osteocytes embedded in the fluid-filled lacuno-canalicular network (LCN). When a bone is loaded, the LCN deforms, causing interstitial fluid flow. Additionally, frequency– and amplitude-dependent pressurization of the intramedullary cavity generates pressure gradients, leading to fluid movement into the marrow space. Varying the dynamic parameters of the load results in distinct responses at both the cellular and tissue levels [3]. Researchers are therefore investigating how specific components of loading, such as frequency, amplitude, duration, fluid velocity, pore pressure and dissipation energy density, affect bone adaptation [4, 5]. Several empirical studies have focused on how cells sense variations in these factors and translates them into new bone formation.

The forces generated by fluid flow are detected by specialized “mechanosensors” present on osteocytes [6]. Mechanical inputs are thus transduced into biochemical signals that encode the information about the stimulus, activating downstream cellular pathways. Osteocytes integrate these signals and coordinate with osteoblasts (bone-forming cells) and osteoclasts (bone-resorbing cells) to reguate bone remodeling.

Early on, it was hypothesized that the complex cellular events following mechanical loading could be modeled as a control system. In a seminal paper, Frost proposed the “mechanostat” as a redumentary framework to explain the dynamics of new bone formation [7]. Mechanical load is first transduced into calcium signalling, which facilitates the transport of transcription factors and co-regulators of osteogenic genes, ultimately driving new bone formation. This inherent feedback in the system naturally suggests the application of control theory – well established in mechanical systems – to the studyof bone mechanobiology.

Several researchers have emphasized the importance of quantifying complex biological phenomena characterized by multiple time scales and non-linearity [8-10]. Developing a mathematical framework not only allows predictions but also facilitates hypothesis generation, particularly in fields like molecular biology where experiments are laborious, expensive and time-consuming. Thus, we propose to construct a control theory-based model to formalize the intricate mechanobiology of bone formation.

To date, many groups have proposed mathematical models that predict BFR spatially across one or multiple cross-sections of bone, especially under exogenous loading protocols. However, there remains a lack of models addressing how BFR evolves temporally under mechanical loading, largely due to challenges in measuring cellular and tissue parameters as time sequences in current experimental systems. Such studies would be valuable for designing exercise regimens and optimizing sports medicine, as well as for understanding the temporal evolution of bone adaptation in clinical contexts. By providing an idealized standard to compare BFR over time, temporal models could also enhance the prognosis and treatment planning for skeletal disorders.

## 2. Methods

### 2.1 Important Factors Influencing the Development of the Model from the Literature – Biological Basis of Fluid Flow and Pore Pressure as Stimuli

In order to develop our model, we first recognized that the dynamics of new bone formation in response to exogenous loading differs between the periosteal and endocortical surfaces. A seminal study [11] demonstrated that the periosteal surface is less sensitive than the endocortical surface in mice of all ages – young, adult and elderly. Intriguingly, this difference is not explained by higher strains within a given surface, since no correlation was observed between strain distribution and bone formation/resorption on the endocortical surface of loaded adult mice. Although the periosteal (Ps) surface exhibits higher average strains, it remains less responsive. Some studies have hypothesized that the proximity of the endocortical (Ec) surface to bone marrow may underlie its greater responsiveness.

This observation led us to question why periosteal bone dynamics differ from endocortical bone dynamics. Could integrins, Ca^2+^signaling, or ATP dynamics explain this discrepancy? We reviewed several studies to investigate further.

For example, deletion of the integrin α5β1 heterodimer in osteocytes resulted in decreased endosteal and trabecular bone formation but no significant change in periosteal bone formation [12]. Another elegant study by Thi et al. [13] showed that calcium signalling responses to hydrodynamic forces were strongest at attachments points on cell processes, and provided direct evidence that integrins αvβ3 play a central role in mechanosensation, particularly fluid flow sensing. Because αvβ3 integrins are more abundant on cell processes, we initially speculated that inhibiting them would reduce periosteal bone formation under mechanical loading. However, experimental evidence revealed that periosteal lamellar bone formation remained unaffected even after αvβ3 inhibition.

This prompted us to consider that multiple mechanosensors may contribute to mechanitransduction and that endocortical bone formation is more sensitive to integrin knockouts. A key study [14] introduced the concept of “osteocyte mechanosomes,” complexes that collectively sense and transduce mechanical loading. These include ATP-gated purigenic receptors (P2X7R), Panx1 channels, CaV3.2-1 voltage-gated channels located near β3 integrin attachments.

Panx1 channels play an important role by releasing ATP in response to loading. Autocrine / paracrine ATP signaling then mediates Ca2+ responses through purinergic (P2) receptors. Indeed, a study showed reduced periosteal mineralizing surface (MS/BS) and bone formation rate (BFR/BS) in loaded Panx1−/− femurs compared to non-loaded Panx1−/− controls [15]. Interestingly, mineral apposition rates on periosteal and endocortical surfaces were similar in wild-type and Panx1−/− mice, regardless of loading, emphasizing ATP signaling’s unique contribution to periosteal mechanotransduction.

Further evidence comes from studies on mitochondrial activity. It was found that periosteal osteocytes exhibit significantly higher active mitochondrial activity, whereas endosteal osteocytes contain more non-functional mitochondria. This could partly explain the differential bone dynamics: periosteal osteocytes may generate more ATP to amplify mechanotransduction, compensating for reduced fluid flow at the periosteal surface due to its impermeability [16], [17].

Several studies have further highlighted the importance of αvβ3 integrins in sensing fluid shear stress (FSS) [18], with mathematical models proposed to capture this phenomenon [19]. Inhibition of αvβ3 integrins attentuated PI3K-AKT activation in response to loading, which in turn blocked α5 activation and hemichannel opening [20]. In vitro studies also showed that blocking these integrins reduced COX-2 expression and PGE2 release under fluid shear stress [21]. Yet, despite their sensitivity to fluid flow, knock out of α5β3 or αvβ3 integrins did not inhibit periosteal bone formation under mechanical loading [6, 22]. Even in mice lacking connexin 43 (Cx43) in osteocytes and osteoblasts, periosteal BFR remained unaffected [23].

Since αvβ3 integrin are primary sensors of fluid flow, these findings suggest that periosteal bone formation may not be driven predominantly by fluid flow, but rather by other physical factors such as pore pressure [24]. This aligns well with our model, which identifies dissipation energy density (DED) – arising from both fluid flow and pore pressure – as the key mechanobiological stimulus and optimization parameter for new bone formation [24].

### 2.2 Osteogenic Signaling Pathway Leading to Type I Collagen – Osteocyte Control and Regulation of Osteogenic Genes: SOST/Sost as a Mediator

Calcium signaling in osteocytes plays a key role in regulating cellular pathways involved in mechanotransduction [25]. As described in [25], osteocyte calcium signaling activates downstream pathways that influence bone remodeling. This signaling within osteocytes further promotes osteoblastogenesis and the up-regulation of osteogenic genes in osteoblasts. Notably, SOST acts as a key negative regulator in this process. The cascade ultimately contributes to the expression of COL1A1, the gene encoding for type I collagen protein, which constitutes about 90% of the organic matrix of bone [26].

In osteoblasts, PIEZO1/2 activates calcium signalling, increasing intracellular Ca^2+^, which in turn stimulates calcineurin. Calcineurin facilitates the nuclear transloacation of β-catenin, thereby enhancing transcription of osteogenic genes. Calcineurin is also involved in the nuclear transport of NFATc1. This process is mediated by Wnt pathway, in which Wnt ligands bind to Frizzled and LRP5/6 receptors on the cell membrane. YAP/TAZ are another downstream elements of PIEZO1/2 signalling; as transcriptional coregulators, they do not bind DNA directly but interact with other transcription factors to amplify their activity in response to mechanical loading. Several studies have indicated their role in skeletal development [27]. Through the complex interplay of these transcription factors, bone cells are able to tightly regulate bone formation processes.

Several studies have shown that Sclerostin (SOST/Sost) plays a central role in mediating the regulatory mechanisms between osteocytes and osteoblasts. Sost is highly expressed in osteocytes and generally inhibits new bone formation; it contributrs to bone resorption. Mechanistically, SOST inhibits Wnt signalling by binding to LRP5/6, thereby preventing activation of the Wnt/β-catenin pathway – a critical driver of osteoblastogenesis and bone formation. Importantly, in vivo mechanical loading has been shown to downregulate Sost gene expression and, consequently, reduce sclerostin protein concentration [28]. Another study demonstrated that Ca^2+^ influx through TRPV4 (transient receptor potential cation channel subfamily V member 4) channels regulates the intracellular levels of sclerostin, whereas Ca^2+^ oscillations specifically influence Sost gene expression [29]. Furthermore, Sost downregulation has been found to be necessary for the anabolic osteogenic response induced by exogenous mechanical loading [30].

Hence, there is a profound connection between calcium signalling in osteocytes, regulation of SOST, and the expression of osteogenic genes in osteoblasts. The Wnt/β-catenin pathway remains central in this network, as it directly promotes the transcription of genes such as COL1A1, thereby driving the production of type I collagen and enabling new bone formation [31]. Together, these insights highlight how mechanical cues are transduced into molecular signals that tightly coordinate bone remodeling.

### 2.3 Relative Gene Expression of Col1a1 as a Simplified Linear Second-Order Critically Damped System

Roosa et al. [26] performed a time-sequenced loading protocol in which Lewis rats were mechanically loaded daily. An axial compressive load was applied as an oscillating Haversine waveform for 360 cycles at a frequency of 2 Hz. The right forelimb bone was loaded axially for 3 minutes per day, while the left forelimb served as the control. The peak load applied was 13 N. The study groups corresponded to durations, measured in days or hours, following the initial application of in vivo bone loading. The rats were loaded for 4 weeks. At designated time intervals, the animals were anesthetized with isoflurane and subsequently euthanized by cervical dislocation.

We assume that the instantaneous bone formation rate (*BFR*(*t*)) induced by external loading is directly proportional to the relative gene expression (*Col*1*a*1(*t*)), but with a time delay *τ*_1_:

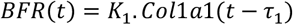

where *K*_1_ is the constant of proportionality.

In the Laplace-domain:

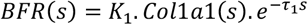

Studies have reported an approximately 96 h delay between loading and the initiation of new bone formation (i.e., observation of non-zero BFR) [32]. We, therefore, modelled the BFR as a linear second-order critically damped system with dead time between application of loading and actual bone formation.

Previous work in our group has shown that bone formation rate is directly proportional to 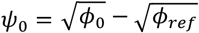, which is square root of the initial dissipation energy (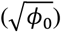) above a threshold or reference value (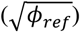) [24]. Since the experimental loading protocol remains constant throughout, we can take *ψ*_0_ as a step function equal to a constant value applied every day (2 Hz Haversine waveform with a peak load of 13 N for 360 loading cycles). This allows us to write:

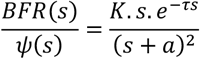

where *K, a* and *τ* are constants to be determined by fitting experimental data, and

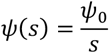

Therefore,

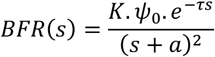

Applying the inverse Laplace transform:

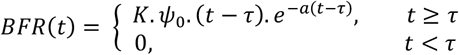

The Col1a1 gene expression is thus given by:

In the Laplace domain:

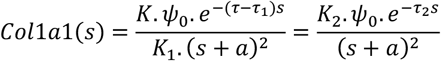

In the time domain:

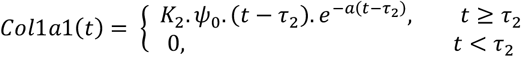

where *K*_2_*ψ*_0_, *a* and *τ*_2_ are constants to be determined by fitting experimental data.

### 2.4. Determination of Parameters *K, a* and *τ*

Roosa et al. [26] reported relative Col1a1 gene expression as a function of time. The values of *K*_2_*ψ*_0_, *a* and *τ*_2_ were obtained by fitting our *Col*1*a*1(*t*) model to the experimental data.

The expression for *BFR*(*t*) contains *Kψ*_0_, *a* and *τ*, where *a* is already determined from curve-fitting of *Col*1*a*1(*t*). *τ* may be assumed to be 96 hours, in accordance with the literature [32]. *Kψ*_0_ can be determined by fitting experimental data. However, no time-sequence BFR data are available for rats under the loading protocol used by Roosa et al. [26]. Nevertheless, an average BFR value for the interlabeling period can be obtained for this loading protocol from the study of Hsieh et al. [33].

## 3. Results and Discussion

Using the Levenberg-Marquardt algorithm [34-35] in MATLAB (Mathworks Inc.) for curve fitting to the experimental values reported by Roosa et al. [26], the following parameters were obtained: *K*_2_*ψ*_0_ = 1.6119 per day, *a* = 0.1374 per day, and *τ*_2_ = 3.4153 days. The comparision of the modeled Col1a1 gene expression with the experimental values is shown in Fig. 1.

**Figure 1.**
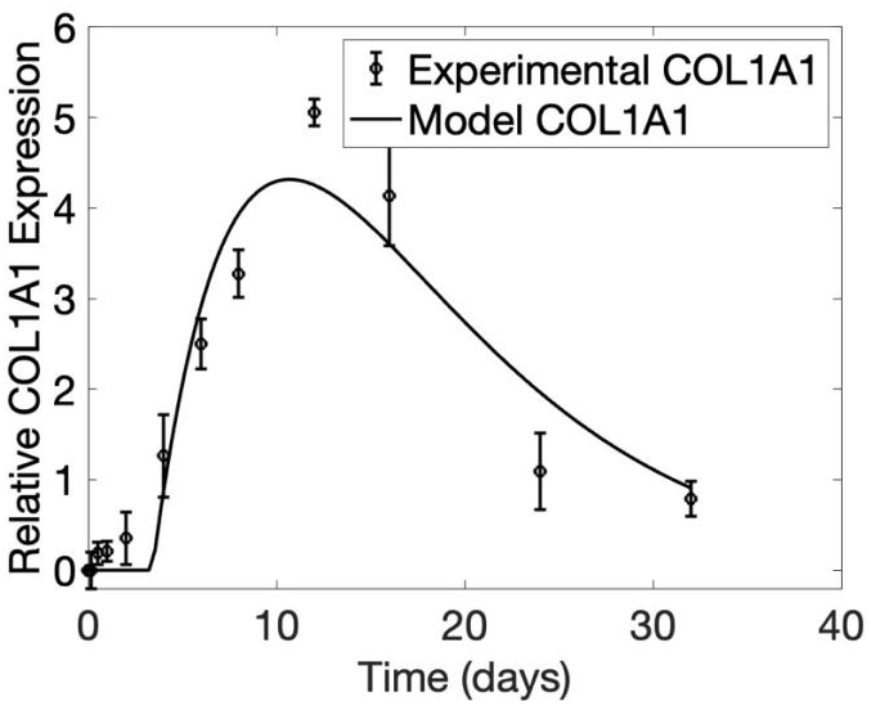
Comparison of relative Col1a1 gene expression: model vs. experimental values

Hsieh et al. [33] reported the average BFR (i.e., 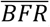) for 10.5N and 14.5N loads, whereas the Col1a1 gene expression data of Roosa et al. [26] correspond to 13N load. By interpolation, we obtained the corresponding average BFR to be 278.4 (standard deviation (SD) = 231) μm/year, i.e., 0.763 (SD = 0.633) μm/day. This value corresponds to *t*_1_ = 5 days, and *t*_2_ = 12 days. By fitting the instantaneous BFR (i.e., *BFR*(*t*)) with *a* = 0.1374 per day and *τ* = 4 days, to the average BFR (i.e., 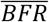), we obtained *Kψ*_0_ = 0.2751 μm/day^2^. The corresponding instantaneous BFR is plotted in Fig. 2.

**Figure 2.**
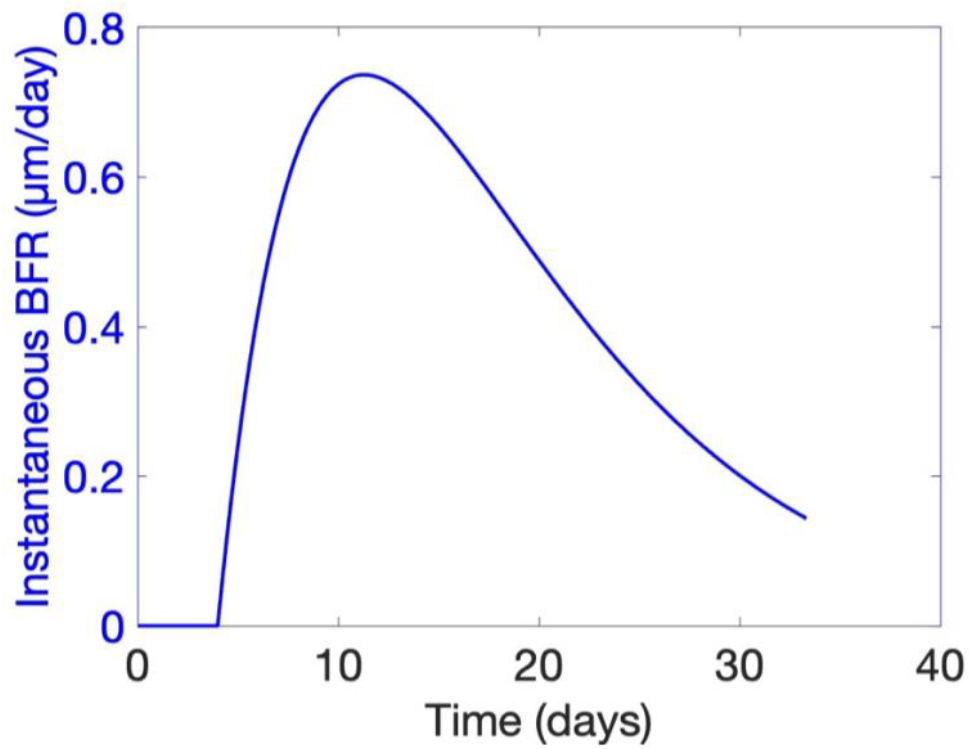
Instantaneous bone formation rate for the loading protocol of Roosa et al. [26].

Both *Col*1*a*1(*t*) and *BFR*(*t*) are shown together in Fig. 3 to compare there their tempral trends. It can be seen that *BFR*(*t*) follows *Col*1*a*1(*t*), with a delay of *τ*_1_ = *τ* − *τ*_2_ = 0.5847 day.

**Figure 3.**
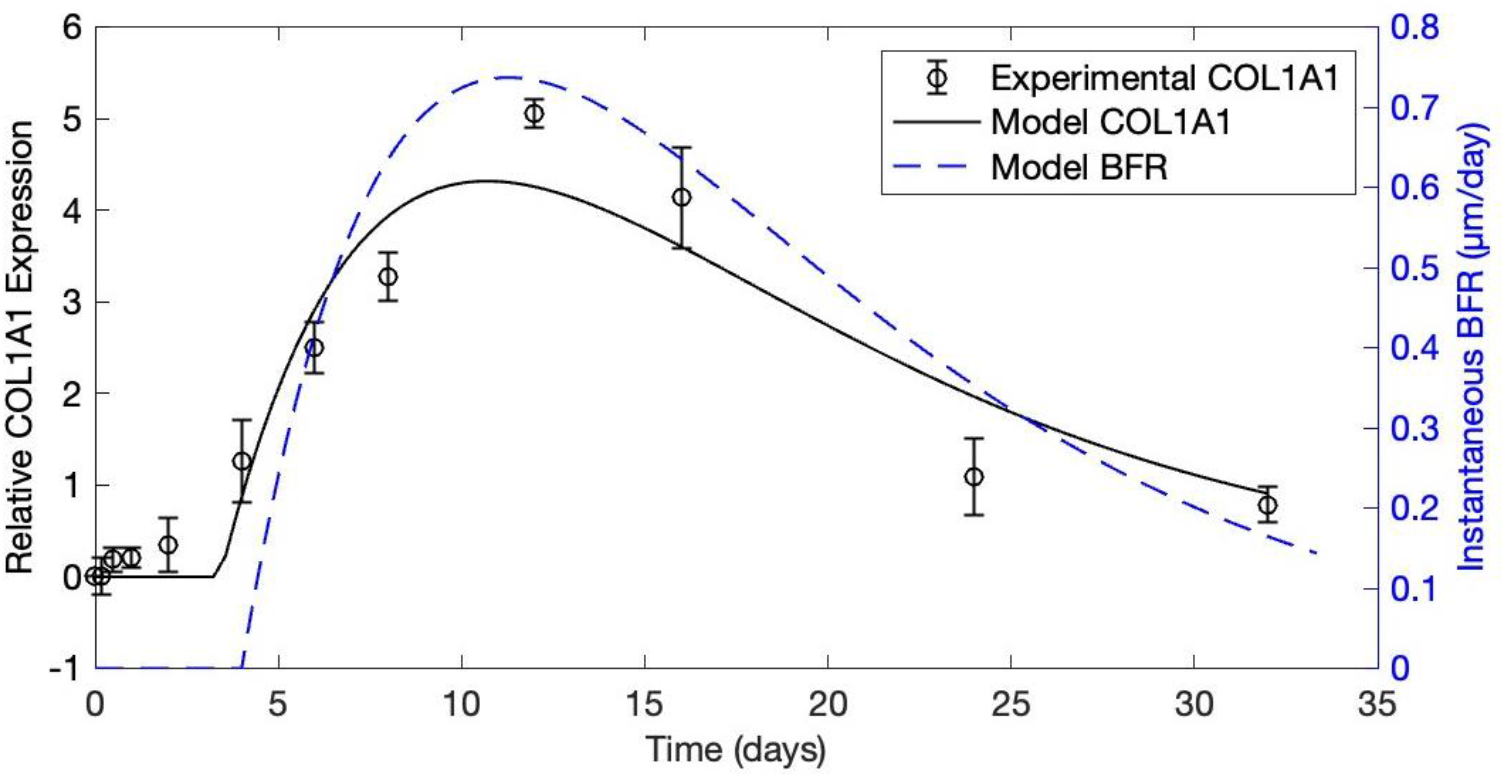
Experimental *Col*1*a*1(*t*), modeled *Col*1*a*1(*t*) and modeled *BFR*(*t*)

The Col1A1 model follows the experimental trend; however, there are significant errors at some time points. Nevertheless, this study demonstrates a methodology to develop a mathematical model to predict instantaneous bone formation rate from temporal data of Col1a1 gene expression. Moreover, this work led to improved results by considering a higher-order model [36]. Subsequently, based on the present work, and Prasad and Aruva [36], a generalized mathematical model was derived by Prasad [37].

## 4. Conclusions and Future Work

This study is among the few that analyze new bone formation temporally in response to exogenous loading. By formalizing the mechanotransduction cascade, we quantified how mechanical loading is encoded within biological parameters. Our findings highlight three key parameters that characterize this encoding:

(i) *τ* – the delay before initiation of new bone formation following mechanical loading,
(ii) *a* – the rate constant influencing the decay of gene expression/BFR, and
(iii) *K* – the proportionality factor reflecting the magnitude of response.

Together, these parameters provide a simplified yet mechanistically grounded framework to describe how bone responds to repetitive loading. The present prototype model, based on a second-order critically damped system with dead time, reproduces experimental observations under a specific loading regimen (2 Hz Haversine waveform, 13 N peak load).

Although the model captures the overall trend of Col1a1 expression and BFR, discrepancies remain at certain time points. These deviations emphasize the need for refinement. Future work could extend this framework by:

(a) Incorporating higher-order or fractional-order dynamics to better capture non-linear biological responses [36-37],
(b) Accounting for additional biomarkers beyond Col1a1 to reflect broader osteogenic signaling,
(c) Integrating variability in experimental conditions (e.g., loading frequency, amplitude, and cycle number) [37], and
(c) Validating predictions against a larger set of in vivo datasets across species and skeletal sites.

By systematically expanding the model, it will be possible to derive a generalized mathematical framework that links cellular biomarkers to instantaneous bone formation rates with greater accuracy. Such advancements hold promise for guiding exercise prescriptions, improving strategies in sports medicine, and informing clinical interventions for skeletal disorders.

### CRediT authorship contribution statement

#### Achsah Marlene Aruva

Literature Review, Conceptualization, Methodology, Investigation, Formal analysis, Data curation, Writing-original draft, review. **J.Prasad**: Conceptualization, Methodology, Investigation, Supervision, Writing-original draft, review, editing.

#### Declaration of Competing Interest

The authors declare that they have no known competing financial interests or personal relationships that could have appeared to influence the work reported in this paper.

## Acknowledgment

The author(s) would like to acknowledge IIT Ropar for providing facilities to carry out the present work.

## Notes

### Competing Interest Statement

The authors have declared no competing interest.

